# Implications Derived from S-Protein Variants of SARS-CoV-2 from Six Continents

**DOI:** 10.1101/2021.05.18.444675

**Authors:** Sk. Sarif Hassan, Kenneth Lundstrom, Pabitra Pal Choudhury, Giorgio Palu, Bruce D. Uhal, Ramesh Kandimalla, Murat Seyran, Amos Lal, Samendra P. Sherchan, Gajendra Kumar Azad, Alaa A. A. Aljabali, Adam M. Brufsky, Ángel Serrano-Aroca, Parise Adadi, Tarek Mohamed Abd El-Aziz, Elrashdy M. Redwan, Kazuo Takayama, Debmalya Barh, Nima Rezaei, Murtaza Tambuwala, Vladimir N. Uversky

**Author notes:** Email addresses:* (Sk. Sarif Hassan), (Kenneth Lundstrom), (Pabitra Pal Choudhury), (Giorgio Palu), (Bruce D. Uhal), (Ramesh Kandimalla), (Murat Seyran), (Amos Lal), (Samendra P. Sherchan), (Gajendra Kumar Azad), (Alaa A. A. Aljabali), (Adam M. Brufsky), (Ángel Serrano-Aroca), (Parise Adadi), (Tarek Mohamed Abd El-Aziz), (Elrashdy M. Redwan), (Kazuo Takayama), (Debmalya Barh), (Nima Rezaei), (Murtaza Tambuwala), (Vladimir N. Uversky).

## Abstract

Spike (S) proteins of severe acute respiratory syndrome coronavirus 2 (SARS-CoV-2) are critical determinants of the infectivity and antigenicity of the virus. Several mutations in the spike protein of SARS-CoV-2 have already been detected, and their effect in immune system evasion and enhanced transmission as a cause of increased morbidity and mortality are being investigated. From pathogenic and epidemiological perspectives, spike proteins are of prime interest to researchers. This study focused on the unique variants of S proteins from six continents Asia, Africa, Europe, Oceania, South America, and North America. In comparison to the other five continents, Africa (29.065%) had the highest percentage of unique S proteins. Notably, only North America had 87% (14046) of the total (16143) specific S proteins available in the NCBI database(across all continents). Based on the amino acid frequency distributions in the S protein variants from all the continents, the phylogenetic relationship implies that unique S proteins from North America were significantly different from those of the other five continents. Overtime, the unique variants originating from North America are most likely to spread to the other geographic locations through international travel or naturally by emerging mutations. Hence it is suggested that restriction of international travel should be considered, and massive vaccination as an utmost measure to combat the spread of COVID-19 pandemic. It is also further suggested that the efficacy of existing vaccines and future vaccine development must be reviewed with careful scrutiny, and if needed, further re-engineered based on requirements dictated by new emerging S protein variants.

## 1. Introduction

The world is experiencing a health emergency due to Coronavirus disease (COVID-19), caused by a deadly enveloped positive-sense single-stranded RNA virus, severe acute respiratory syndrome coronavirus (SARS-CoV-2) [1, 2, 3, 4, 5, 6]. The spike (S) protein is a homotrimer present on the surface of the SARS-CoV-2 and recognizes the human host cell surface receptor angiotensin-converting enzyme-2 (ACE2) [7, 8, 9, 10]. From the beginning of the second wave of COVID-19 infection, various SARS-CoV-2 variants variants emerged raising concern of enhanced transmission and mortality of the virus and reduced efficacy of vaccine protection [11, 12]. Some of the studies opposed the perception of SARS-CoV-2 mutations as distinctive pathogenic variants and increased rate of transmissibility were questioned [13, 14]. However, the frequency of the mutant strains within the SARS-CoV-2 population carrying the D614G mutation in the spike protein clearly plays a role in enabling the virus to spread more effectively and rapidly [15]. Epidemiologists have been constantly monitoring the evolution of SARS-CoV-2 with a particular focus on the spike protein and other interacting proteins of the virus [15, 16]. The D614G mutation in the S protein discovered in early 2020 makes the virus able to spread more effectively and rapidly [17]. The D614G mutation has been found to be related with high viral loads in infected patients, and high rate of infections, but not with increased disease severity [18]. Various mutations in the S protein make the SARS-CoV-2 more complex and hence it is more difficult to characterize its severity, infectivity and efficacy of vaccines designed to target S protein. Not all mutations are advantageous to the virus but several mutations or a set of mutations may increase the transmission potential through an increase in receptor binding or the ability to evade the host immune response by altering the surface structures recognized by antibodies [19, 20, 21].

To contain the spread of the COVID-19, it is definitely of high interest to detect and identify various unique emerging variants of S proteins. Additionally, it is also worth investigating the impact of new S protein variants on viral infectivity and potential to spread rapidly as well as to acertain the origin of the spread of the new variants concerning spike protein variabilities. Accordingly, it might be possible to segregate the set of new variants with respect to individual characteristics of SARS-CoV-2, which would undoubtedly help policy makers to form various strategies to contain the spread of the virus. There are a large number of different SARS-CoV-2 S protein mutant sequences currently available in the NCBI virus database. In this study, all available S protein sequences from six continents Asia, Africa, Europe, North America, South America, and Oceania were analyzed for their uniqueness and variability. An inter-linkage was made among the unique S proteins available on the six continents was performed.

## 2. Data acquisition and methods

S protein sequences from all six continents (Asia, Africa, Europe, Oceania, South America, and North America) were downloaded in Fasta format (on May 7, 2021) from the National Center for Biotechnology Information (NCBI) database (http://www.ncbi.nlm.nih.gov/). Further, fasta files were processed in *Matlab-2021a* for extracting unique S protein sequences for each continent.

### 2.1. Frequency probability of amino acids

Any protein sequence is composed of twenty different amino acids with various frequencies starting from zero. The probability of occurrence of each amino acid *A_i_* is determined by the formula 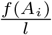 where *f*(*A_i_*) denotes the frequency of occurrence of the amino acid *A_i_* in a primary sequence, and *l* stands as the length of an S protein [22]. Hence for each S protein, a twenty-dimensional vector considering the frequency probability of twenty amino acids can be obtained. Based on this frequency probability, the dominance of amino acid density in a given protein is illuminated.

### 2.2. Evaluation of normalized amino acid compositions

The variability of the amino acid compositions of the unique S-proteins from each continent was evaluated using the webbased tool Composition Profiler (http://www.cprofiler.org/) that automates detection of enrichment or depletion patterns of individual amino acids or groups of amino acids in query proteins [23]. In this analysis, we used sets of unique S-proteins from each continent as query samples and the amino acid of the original S-protein (UniProt ID: P0DTC2) as a reference sample that provides the background amino acid distribution. Composition profiler generates a bar chart composed of twenty data points (one for each amino acid), where bar heights indicate normalized enrichment or depletion of a given residue. The normalized enrichment/depletion is calculated as

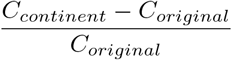

where *C_continent_* is the content of given residue in the query set of S-proteins in a given continent and *C_original_* is the content of the same residue in the original S-protein. For comparison, we generated composition profile of disordered proteins, where normalized composition was evaluated as 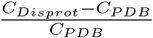 (*C_DisProt_* = content of a given amino acid in the set of intrinsically disordered proteins in the DisProt database [24]; *C_PDB_* = content of the given residue in the dataset of fully ordered proteins, PDB Select 25 [23]). In these analyses, the positive and negative values produced in the compositional profiler indicated enrichment or depletion of the indicated residue, respectively.

### 2.3. Amino acid conservation Shannon entropy

How conserved/disordered the amino acids are organized over S protein is addressed by the information-theoretic measure known as *‘Shannon, entropy*(SE)’. For each S protein, Shannon entropy of amino acid conservation over the amino acid sequence of S protein is computed using the following formula [25, 26]:

For a given amino acid sequence of length *l*, the conservation of amino acids is calculated as follows:

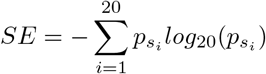

where 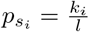; *k_i_* represents the number of occurrences of an amino acid *s_i_* in the given sequence [27].

### 2.4. Isoelectric point of a protein sequence

The isoelectric point (IP), is the pH at which a molecule carries no net electrical charge or is electrically neutral in the statistical mean. We calculate the theoretical pI by using the pKa’s of amino acids and summing the net charge across the protein at a given pH (default is typical intracellular pH 7.2), searching with our algorithm for the pH at which the net charge is zero [28].

Note that the isoelectric point of a protein sequence was computed here using the standard routine of *Matlab-2021a*. This parameter was deployed to characterize the unique S protein sequences, quantitatively.

## 3. Results

We first determined the set of unique S protein sequences from each continent. Further, every unique S protein from a continent was compared with other unique S proteins from five continents, and the lists of the same are presented in Tables 12–17. Also, the variability of the S proteins from each continent was shown using Shannon entropy and isoelectric point.

### 3.1. Unique spike proteins in the continents

In Table 1, the number of total sequences, unique sequences and percentages are presented. Note that, a complete list of unique S protein accessions and their names (continent-wise) were made available in *supplementary file-1*. Note that, sequence accession is renamed as *Ck* where *C* stands for continent code (Asia:AS, Africa:AF, Oceania:O, Europe:U, South America:SA and North America:NA), and *k* denotes the serial number.

**Table 1:** Percentages of continent-wise unique spike (S) proteins

| Continent | Total S proteins (T) | Unique S proteins (U) | Percentage, continent-wise<br>$\left(\frac{U}{T} \times 100\right)$ | Percentage, worldwide<br>$\left(\frac{U}{16143} \times 100\right)$ |
| --- | --- | --- | --- | --- |
| *Africa* | 984 | 286 | 29.065 | 1.772 |
| *Asia* | 2314 | 432 | 18.669 | 2.676 |
| *Europe* | 1006 | 187 | 18.588 | 1.158 |
| *Oceania* | 9920 | 1121 | 11.300 | 6.944 |
| *South America* | 464 | 71 | 15.302 | 0.440 |
| *North America* | 113072 | 14046 | 12.422 | 87.010 |
| *Worldwide* | 127760 | 16143 | 12.635 | — |

The highest amount (29.065%) of unique S proteins were found in Africa though the total number of available sequences is significantly low as compared with that from other continents. Almost similar amounts (in percentage) of unique S sequence variations were found in Asia and Europe. Among the total 127760 S proteins embedded in SARS-CoV-2 genomes, only 16143 (12%) unique S proteins were detected so far, and notably most of the unique variants (87%) were found in North America only.

For each continent, the unique spike (S) proteins were matched with other unique proteins from the rest of the five continents, and a total number of such identical pairs are presented accordingly in the matrix (Table 2).

**Table 2:** Continent-based frequencies of identical S proteins

| Continent-wise | Asia | Africa | Europe | North America | Oceania | South America |
| --- | --- | --- | --- | --- | --- | --- |
| *Asia* | — | 25 | 27 | 169 | 17 | 17 |
| *Africa* | 25 | — | 15 | 71 | 13 | 5 |
| *Europe* | 27 | 15 | — | 76 | 9 | 8 |
| *North America* | 169 | 71 | 76 | — | 49 | 31 |
| *Oceania* | 17 | 13 | 9 | 49 | — | 5 |
| *South America* | 17 | 5 | 8 | 31 | 5 | — |
| *Total continent-wise* | 255 | 129 | 135 | 396 | 93 | 66 |
| *Unique residue S proteins* | 177 | 157 | 52 | 13650 | 1028 | 5 |

From Table 2, it was observed that, in each continent there is still a significant percentage of unique spike variations available, which are not shared with any rest of the continents. Such percentages of unique variations of S proteins in Asia, Africa, Europe, Oceania, South America, and North America were 41%, 55%, 28%, 92%, 7%, and 97% respectively. The lists of pairs of identical S proteins of SARS-CoV-2 originating from six continents are presented in Tables 9–11 (*Appendix*-I).

The lists of unique S proteins (from a particular continent), which were found to be identical with some unique spike proteins from other five continents, are presented in Tables (12–17) (*Appendix*-II).

The frequency and percentage of invariant residue positions, where no amino acid change was detected so far in the unique S proteins available in each continent, are presented in Table 3.

**Table 3:** Frequency and percentage of invariant residue positions among 1273 positions in unique S proteins

|  |  |  |  |  |  |  |
| --- | --- | --- | --- | --- | --- | --- |
| *Frequency of invariant residue positions in unique S proteins from each continent* |  |  |  |  |  |  |
|  | **Africa** | **Asia** | **Europe** | **Oceania** | **South America** | **North America** |
| **Total Freq.** | 902 | 695 | 948 | 731 | 1070 | 89 |
| **Percentage** | 70.86 | 54.60 | 74.47 | 57.42 | 84.05 | 6.99 |

The highest number of mutations (lowest number of invariant residue position, 6.99%) (Table 3) were detected in the unique S proteins from North America where 12.42% unique S protein sequences were present as mentioned in Table 1. Likewise, the lowest number (15.95%) of mutations in unique S proteins were observed in South America where 15.3% unique S sequences were found. Only 29.14% residues of 1273 in the unique S proteins were mutated, although a significantly higher number (29.065%) of unique sequences were found in Africa among the other five continents. The unique S proteins from Europe possessed only 25.5% mutations, whereas 45.5% mutations were detected in the unique S proteins from Asia although the same percentage (18.5%) of unique spike proteins were found (Table 1 and 3). Further it was observed that 11.3% of the unique S proteins from Oceania possessed 42.58% mutations.

### 3.2. Variability through normalized amino acid composition

Additional information on the variability of the amino compositions of the unique S-proteins from each continent relative to the composition of original S-protein from Wuhan was retrieved using the web-based tool Composition Profiler (http://www.cprofiler.org/). Results of this analysis are shown in Figure 4A, which clearly shows the presence of some noticeable amino acid composition variability among unique S-proteins from different continents. Since individual S proteins are different from each other and from the original S-protein mostly in very limited number of residues, the range of changes in the normalized enrichment/depletion of a given residue is rather limited (compare scales of Y axis in Figures 1A and 1B, where a composition profile of the intrinsically disordered proteins is shown for comparison).

**Figure 1:**
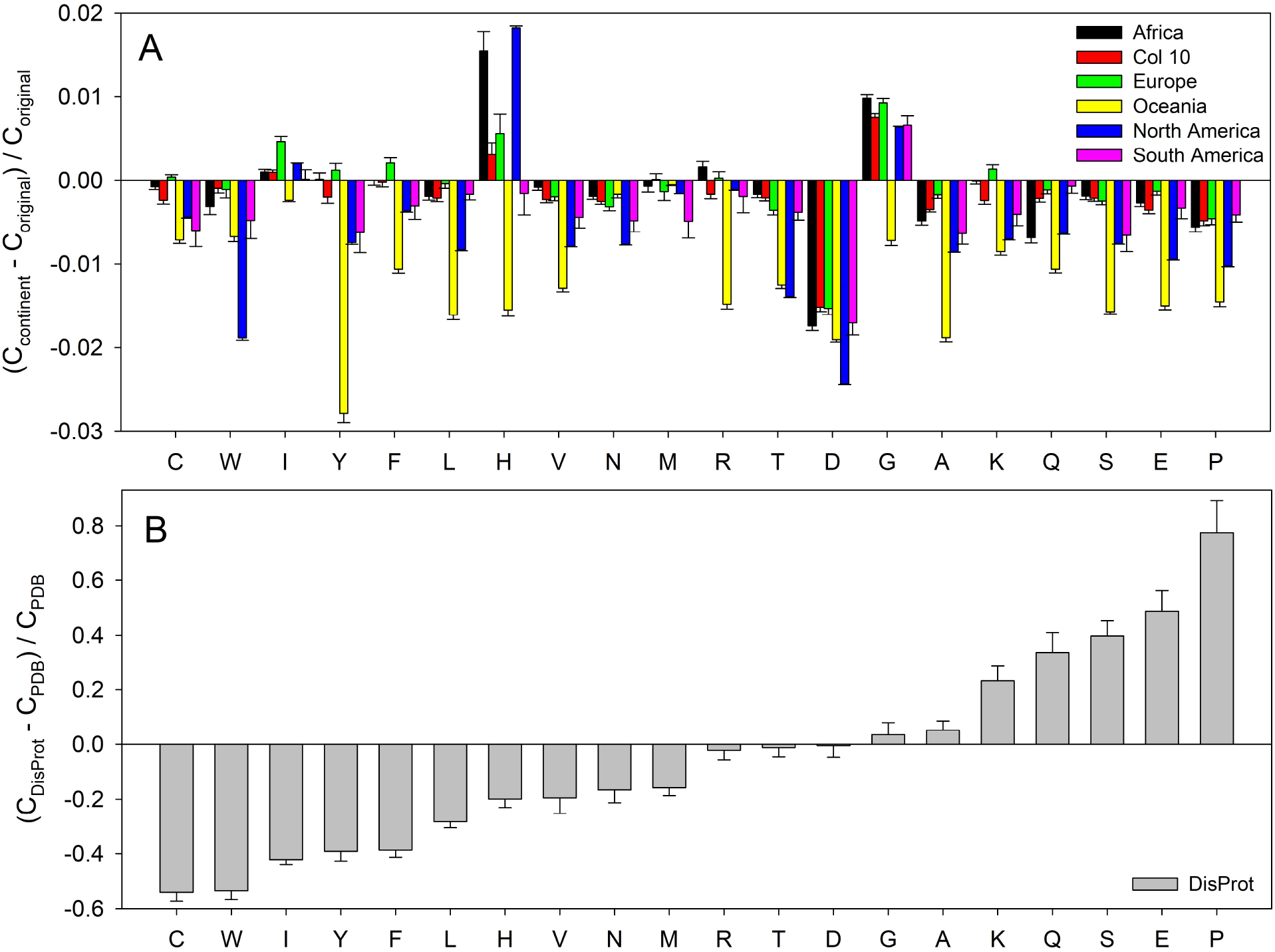
Composition profiles of unique S-proteins from different continents (A) in comparison with the composition profile of typical intrinsically disordered protein (B).

On an average, unique S-proteins form Oceania were found to have the most variability in terms of normalized amino acid composition. This was followed by the unique S-proteins from North America. Curiously, Figure 1A shows that although the normalized content of individual residues in the unique S-proteins from Oceania is always below that of the original S-protein, S-proteins from other continents might have relative excess of some residues. For example, some unique S-proteins from almost all continents can be enriched in glycine or histidine residues, whereas some European S-proteins can also be relatively enriched in cysteine, isoleucine, tyrosine, phenylalanine, and lysine residues (see positive green bars in Figure 1A). Another interesting observation is that the different sets of S-proteins are typically characterized by rather noticeable variability of the normalized content of most residues. The noticeable exception is given by aspartate, depletion in which is almost uniform between all the unique S-proteins from all the continents.

### 3.3. Variability of unique spike proteins

We quantitatively determined the variations in the unique S proteins on six continents. The variations were captured through the frequency distribution of amino acids present, Shannon entropy (amount of conservation of amino acids in a given sequence), and molecular weights and isoelectric points of a given protein sequence.

#### 3.3.1. Variations in the frequency distribution of amino acids

The frequency of each amino acid was computed for each unique S protein available in six continents (*Supplementary file-2*). Maximum and minimum frequencies of amino acids present in the unique S proteins from different continents are presented in Table 4.

**Table 4:** Maximum and minimum frequencies of amino acids present in the unique spike proteins from different continents

| *Max and Min of Frequencies* |  | **A** | **R** | **N** | **D** | **C** | **Q** | **E** | **G** | **H** | **I** | **L** | **K** | **M** | **F** | **P** | **S** | **T** | **W** | **Y** | **V** |
| --- | --- | --- | --- | --- | --- | --- | --- | --- | --- | --- | --- | --- | --- | --- | --- | --- | --- | --- | --- | --- | --- |
| **Africa** | *Max* | 80 | 44 | 89 | 62 | 41 | 63 | 49 | 84 | 19 | 79 | 109 | 62 | 15 | 78 | 60 | 101 | 98 | 13 | 56 | 98 |
|  | *Min* | 73 | 40 | 85 | 58 | 38 | 59 | 45 | 78 | 14 | 73 | 102 | 57 | 13 | 72 | 55 | 94 | 90 | 11 | 49 | 93 |
| **Asia** | *Max* | 80 | 44 | 89 | 63 | 41 | 63 | 49 | 84 | 19 | 78 | 110 | 62 | 15 | 79 | 59 | 101 | 101 | 13 | 57 | 98 |
|  | *Min* | 73 | 39 | 80 | 55 | 36 | 56 | 45 | 76 | 15 | 72 | 100 | 55 | 13 | 68 | 52 | 90 | 90 | 11 | 49 | 90 |
| **Europe** | *Max* | 80 | 43 | 89 | 63 | 41 | 63 | 49 | 84 | 19 | 79 | 110 | 62 | 15 | 79 | 59 | 101 | 98 | 13 | 57 | 99 |
|  | *Min* | 75 | 38 | 84 | 59 | 39 | 59 | 46 | 79 | 16 | 74 | 102 | 58 | 13 | 74 | 54 | 96 | 90 | 11 | 50 | 93 |
| **Oceania** | *Max* | 81 | 43 | 90 | 62 | 41 | 63 | 49 | 84 | 18 | 78 | 109 | 62 | 15 | 79 | 59 | 100 | 98 | 12 | 56 | 99 |
|  | *Min* | 72 | 37 | 81 | 58 | 36 | 57 | 44 | 74 | 15 | 71 | 97 | 56 | 13 | 71 | 52 | 92 | 88 | 10 | 43 | 89 |
| **North America** | *Max* | 82 | 44 | 91 | 63 | 42 | 64 | 49 | 85 | 20 | 79 | 111 | 64 | 15 | 80 | 60 | 102 | 99 | 13 | 58 | 100 |
|  | *Min* | 60 | 32 | 63 | 46 | 32 | 39 | 34 | 63 | 11 | 55 | 82 | 43 | 9 | 55 | 43 | 76 | 77 | 8 | 36 | 82 |
| **South America** | *Max* | 80 | 43 | 89 | 62 | 41 | 63 | 48 | 83 | 18 | 78 | 109 | 62 | 14 | 79 | 58 | 101 | 98 | 12 | 57 | 98 |
|  | *Min* | 75 | 38 | 82 | 57 | 37 | 59 | 45 | 79 | 16 | 73 | 105 | 57 | 13 | 73 | 57 | 92 | 93 | 11 | 50 | 92 |

All S protein sequences are leucine (L) and serine (S) rich. Tryptophan (W) and methionine (M) were presented with the least frequencies (Table 4). The widest variation in frequency distributions of the twenty amino acids over the unique S proteins was found in North America.

To obtain quantitative variations in the unique S proteins available in each continent, differences between maximum and minimum vectors (20 dimensions) were obtained (Table 5), and then Euclidean distances between the difference vectors was calculated (Table 6).

**Table 5:** Matrix presenting the difference between maximum and minimum frequencies of amino acids present in the unique S proteins on each continent

| *Difference matrix* | **A** | **R** | **N** | **D** | **C** | **Q** | **E** | **G** | **H** | **I** | **L** | **K** | **M** | **F** | **P** | **S** | **T** | **W** | **Y** | **V** |
| --- | --- | --- | --- | --- | --- | --- | --- | --- | --- | --- | --- | --- | --- | --- | --- | --- | --- | --- | --- | --- |
| **Africa** | 7 | 4 | 4 | 4 | 3 | 4 | 4 | 6 | 5 | 6 | 7 | 5 | 2 | 6 | 5 | 7 | 8 | 2 | 7 | 5 |
| **Asia** | 7 | 5 | 9 | 8 | 5 | 7 | 4 | 8 | 4 | 6 | 10 | 7 | 2 | 11 | 7 | 11 | 11 | 2 | 8 | 8 |
| **Europe** | 5 | 5 | 5 | 4 | 2 | 4 | 3 | 5 | 3 | 5 | 8 | 4 | 2 | 5 | 5 | 5 | 8 | 2 | 7 | 6 |
| **Oceania** | 9 | 6 | 9 | 4 | 5 | 6 | 5 | 10 | 3 | 7 | 12 | 6 | 2 | 8 | 7 | 8 | 10 | 2 | 13 | 10 |
| **South America** | 5 | 5 | 7 | 5 | 4 | 4 | 3 | 4 | 2 | 5 | 4 | 5 | 1 | 6 | 1 | 9 | 5 | 1 | 7 | 6 |
| **North America** | 22 | 12 | 28 | 17 | 10 | 25 | 15 | 22 | 9 | 24 | 29 | 21 | 6 | 25 | 17 | 26 | 22 | 5 | 22 | 18 |

**Table 6:**
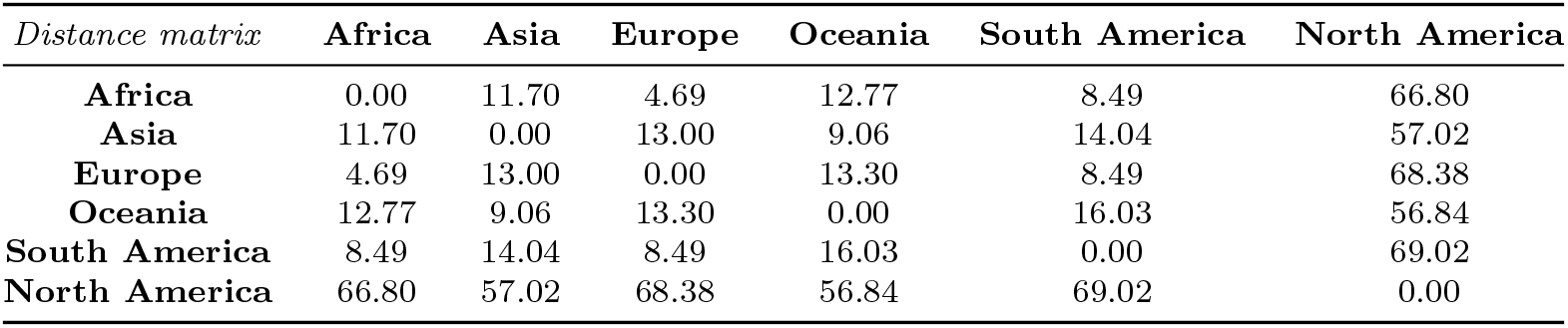
Pairwise Euclidean distances among the difference vectors of each continent

| *Distance matrix* | **Africa** | **Asia** | **Europe** | **Oceania** | **South America** | **North America** |
| --- | --- | --- | --- | --- | --- | --- |
| **Africa** | 0.00 | 11.70 | 4.69 | 12.77 | 8.49 | 66.80 |
| **Asia** | 11.70 | 0.00 | 13.00 | 9.06 | 14.04 | 57.02 |
| **Europe** | 4.69 | 13.00 | 0.00 | 13.30 | 8.49 | 68.38 |
| **Oceania** | 12.77 | 9.06 | 13.30 | 0.00 | 16.03 | 56.84 |
| **South America** | 8.49 | 14.04 | 8.49 | 16.03 | 0.00 | 69.02 |
| **North America** | 66.80 | 57.02 | 68.38 | 56.84 | 69.02 | 0.00 |

**Table 7:** Interval of Shannon entropy of unique S proteins from six different continents

| SE: Continent | Interval of SEs |
| --- | --- |
| SE of S protein: Africa | (0.960825, 0.963239) |
| SE of S protein: Asia | (0.961471, 0.963326) |
| SE of S protein: Europe | (0.961539, 0.963254) |
| SE of S protein: North America | (0.95934, 0.964314) |
| SE of S protein: Oceania | (0.961525, 0.963042) |
| SE of S protein: South America | (0.961589, 0.962895) |

Based on the distance matrix, a phylogenetic relationship was derived among the continents (Figure 2).

**Figure 2:**
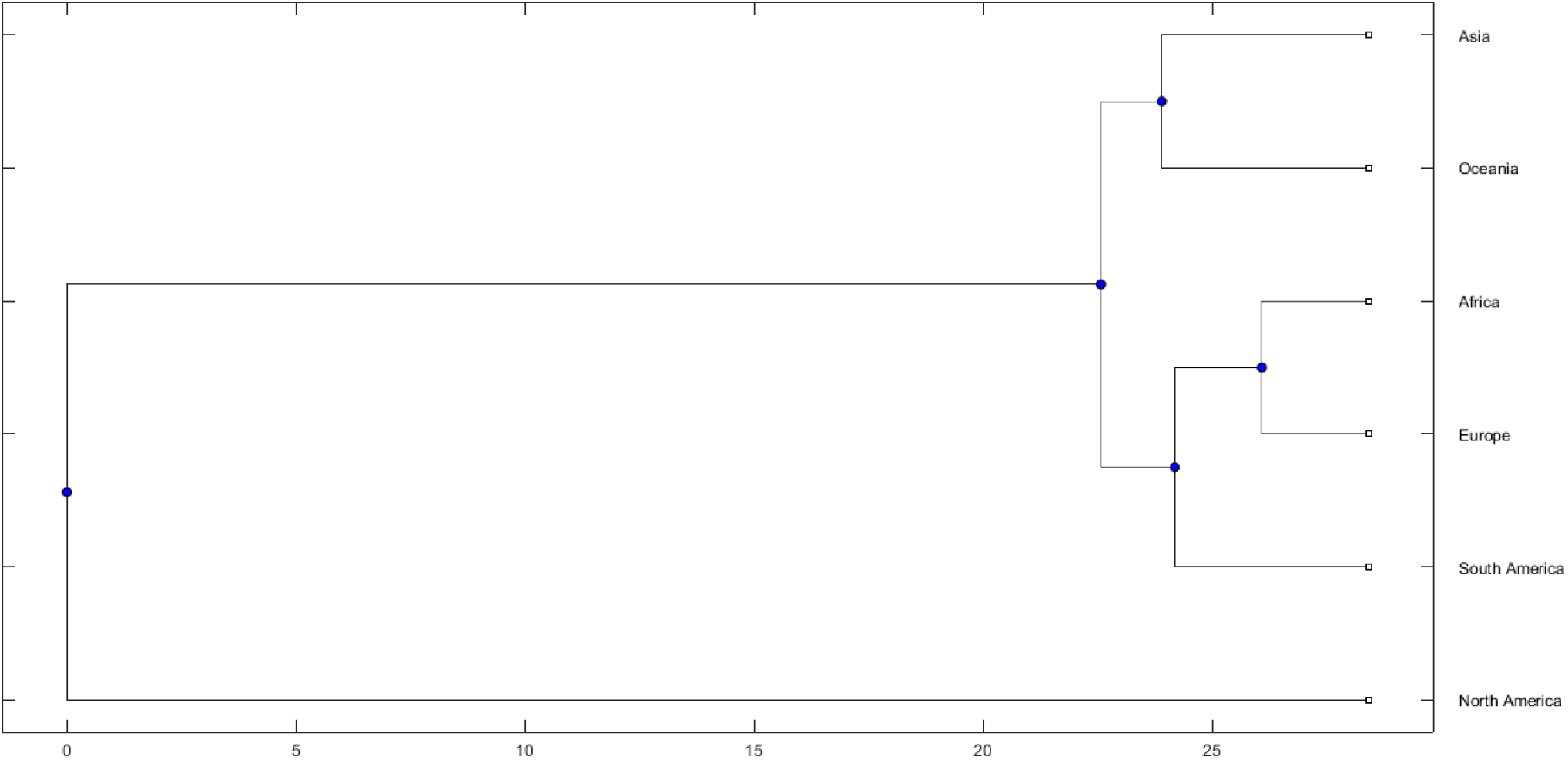
Phylogenetic relationship among the six continents based on the variability of unique spike proteins available in each continent.

Variations based on the frequency distribution of amino acids present in the S proteins make North America (which belongs to the rightmost branch of the tree) distant from the other five continents (Figure 2). Variations among the unique spike proteins from Asia and Oceania turned out to be similar, and they belong to the same level of leaves of the far left branch of the tree. Africa and Europe were found to be the closest in terms of variations based on the frequency distribution of amino acids over the unique spike proteins from each continent. Variability of spike proteins from South America has distant resemblance to that of Africa/Europe as estimated in the phylogeny. The frequencies of amino acid distribution in each unique S protein from each continent were plotted (Figure 3).

**Figure 3:**
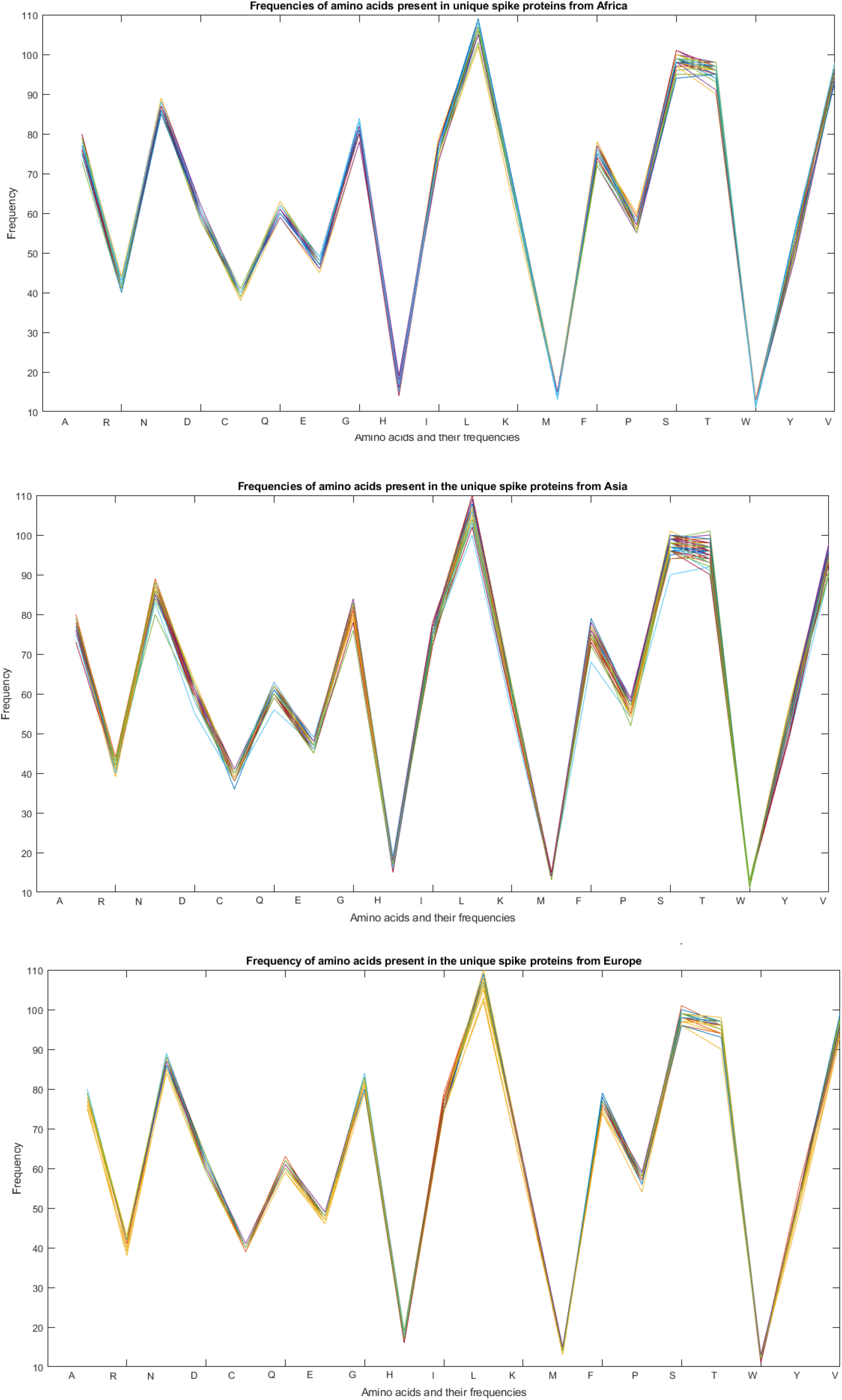
Frequencies of amino acids present in the unique S sequences

**Figure 4:**
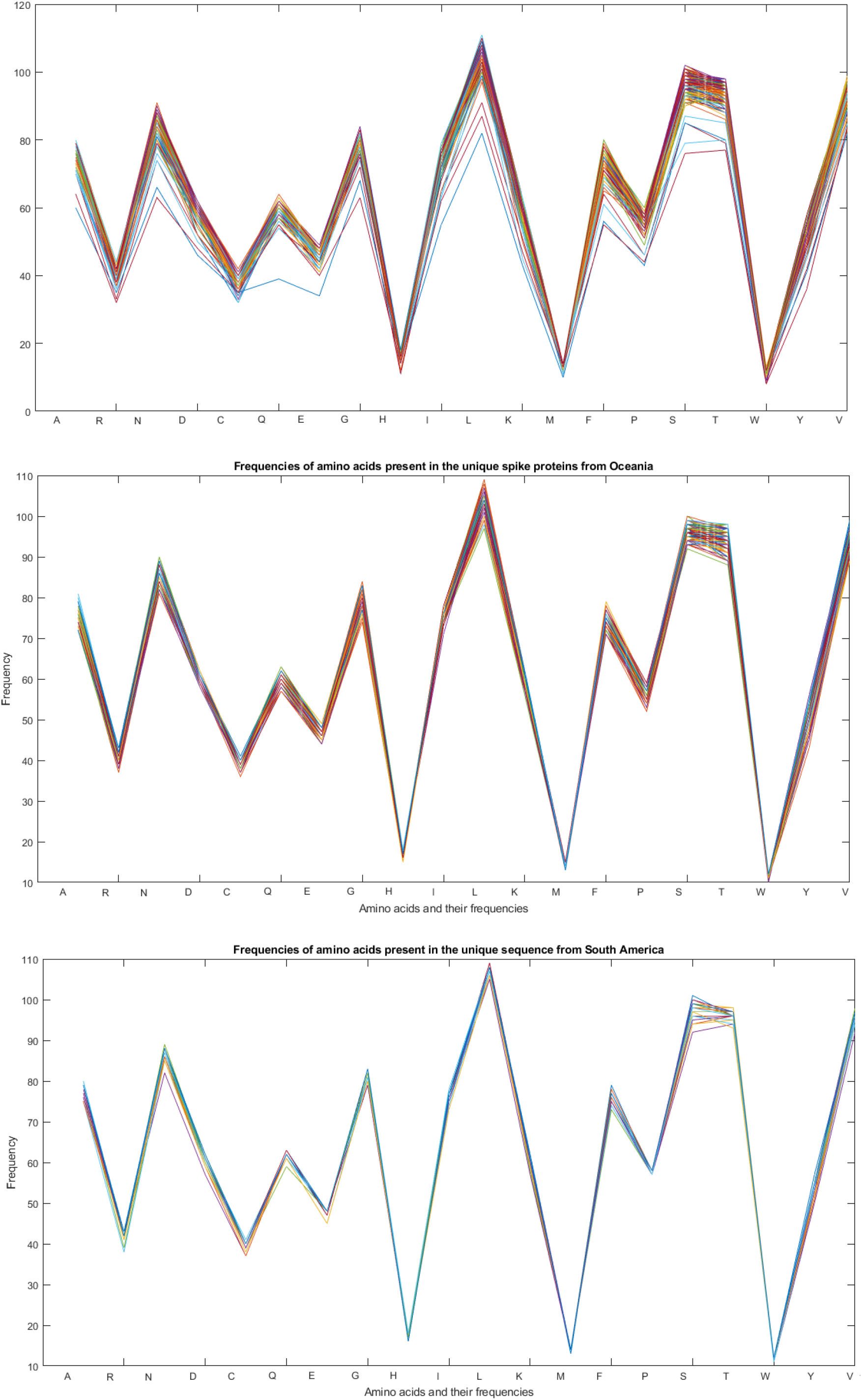
Frequencies of amino acids present in the unique S sequences

The widest variations of the frequency distribution of amino acids present in S proteins were observed in North America as wide band was observed in Figure 4. Individual frequency distributions of amino acids in Asia and Oceania seem very close as it was observed from the phylogeny (Figure 2).

#### 3.3.2. Variability through Shannon entropy

In principle, for a random amino acid sequence, the Shannon entropy (SE) is one. Here Shannon entropy for each S protein sequence was computed using the formula stated in section 2.2 (*Supplementary file-2*). It was found that the highest and lowest SEs of S proteins from all continents were 0.9643 and 0.9594 respectively. That is, the length of the largest interval is 0.005 which is sufficiently small. Also note that the length of the smallest interval was 0.001 which occurred in the SEs of S proteins from South America. Within this realm, the widest variation of SEs was noticed among the unique S proteins of North America. All other four intervals (considering lowest and highest) of SEs of all the unique S proteins from four continents Africa, Asia, Oceania and Europe were contained in the interval of North America and contain that of South America.

Among all (20^1273^) possible amino acids (20 in number) sequences of length 1273, Nature(?) had selected only a fraction to make S proteins of SARS-CoV-2, and interestingly SEs of them were kept within a very small interval. From the SEs which were close to 1, the S protein sequences are expected to be pseudo-random. Variation of SEs for all unique S proteins from each continent is shown in Figures 5 and 6.

**Figure 5:**
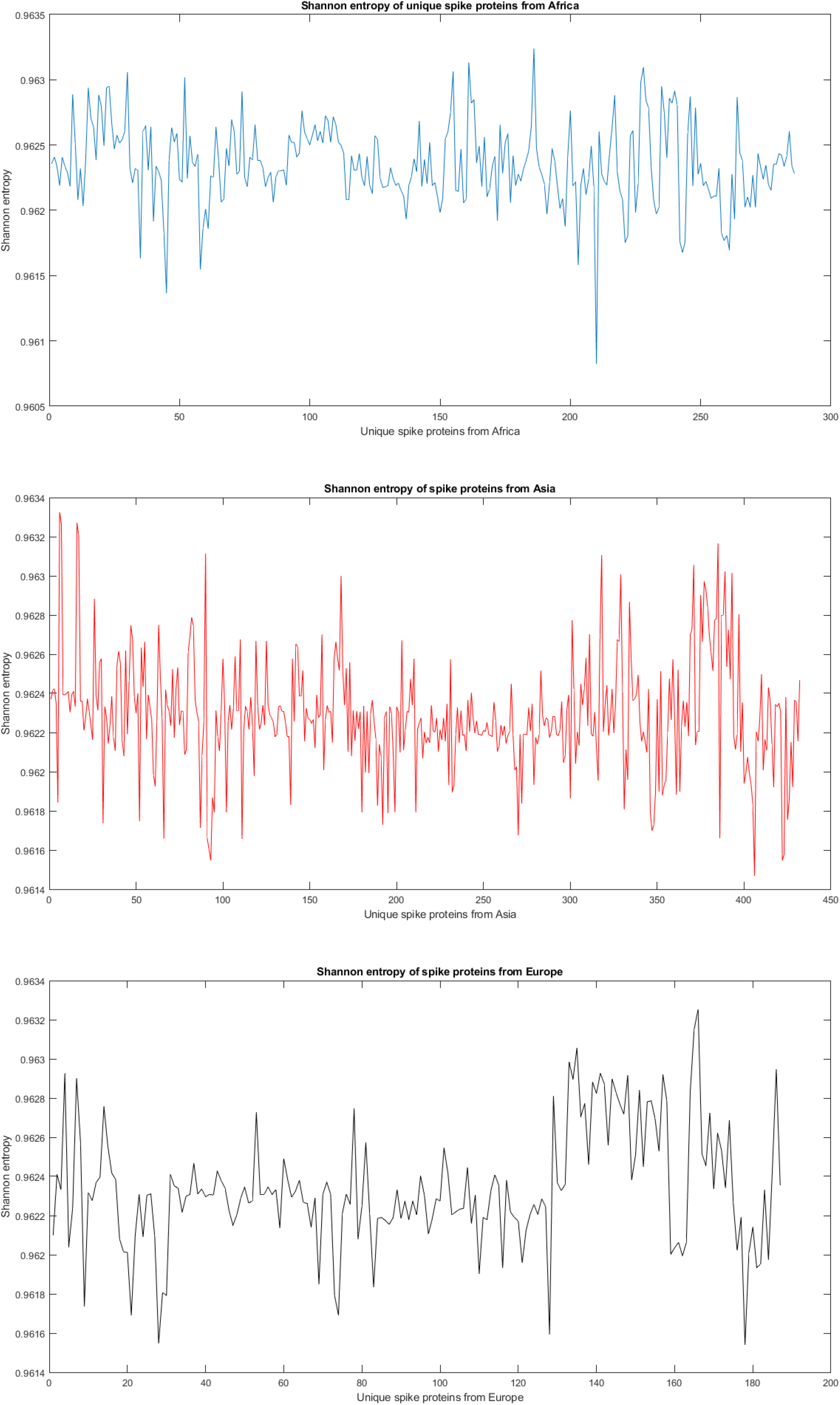
SE of unique S proteins from different continents

**Figure 6:**
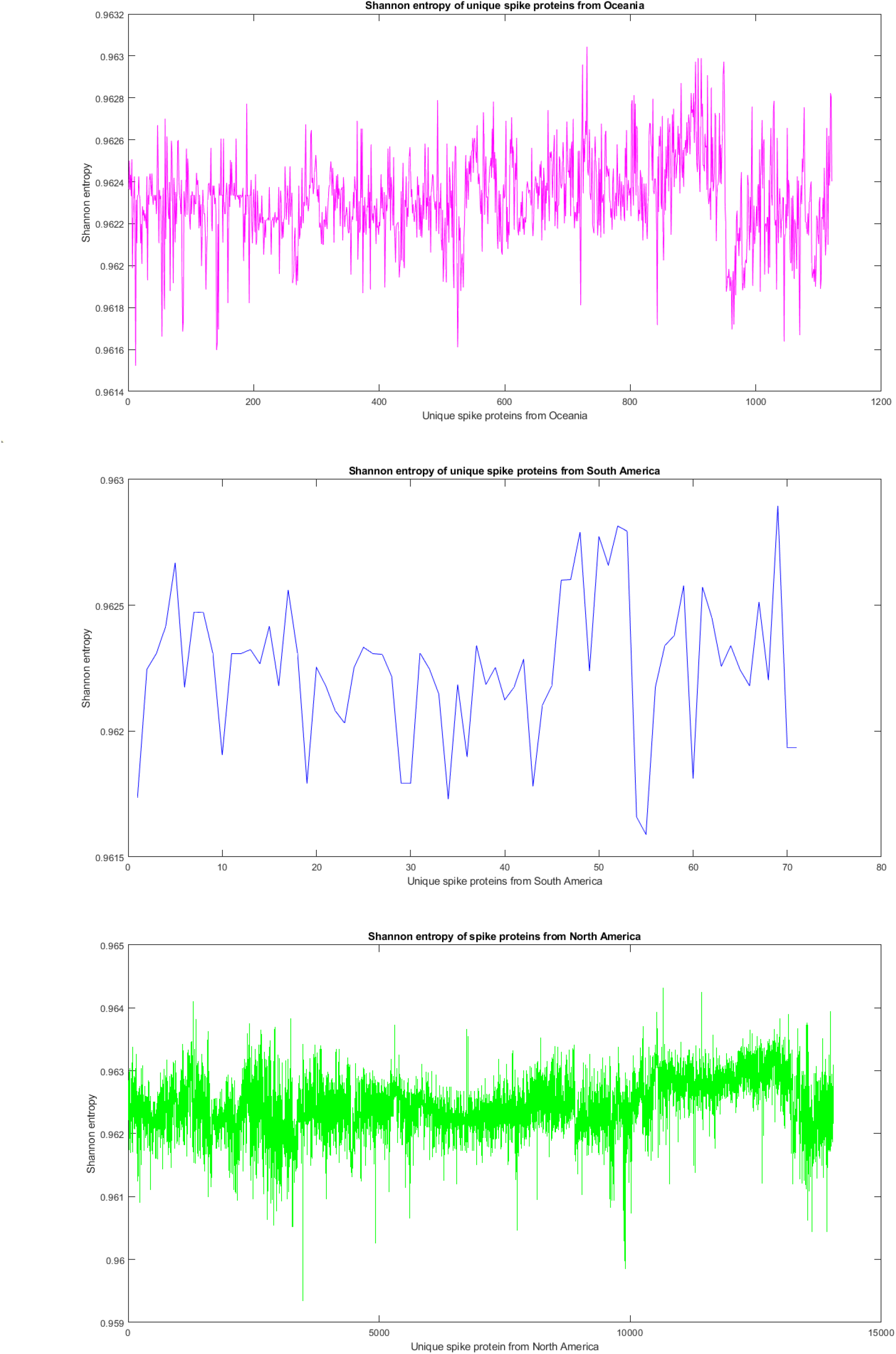
SE of unique S proteins from different continents

Conservation of amino acids present over each S protein from each continent is different from one another which is depicted by the zig-zag nature of SEs plots (Figure 5 and 6).

#### 3.3.3. Variability through isoelectric point

For each S protein sequence from each continent isoelectric point (IP) was computed (Supplementary file-3). Intervals (considering minimum and maximum) IPs of unique spike proteins from each continent were tabulated in Table 8.

**Table 8:** Interval of isoelectric point of unique S proteins from six different continents

| **IP: Continent** | **Interval of IPs** |
| --- | --- |
| IP of S protein: Africa | (6.44, 7.09) |
| IP of S protein: Asia | (6.21, 7.08) |
| IP of S protein: Europe | (6.21, 6.99) |
| IP of S protein: North America | (5.61, 7.79) |
| IP of S protein: Oceania | (6.31, 7.09) |
| IP of S protein: South America | (6.36, 6.99) |

**Table 9:**
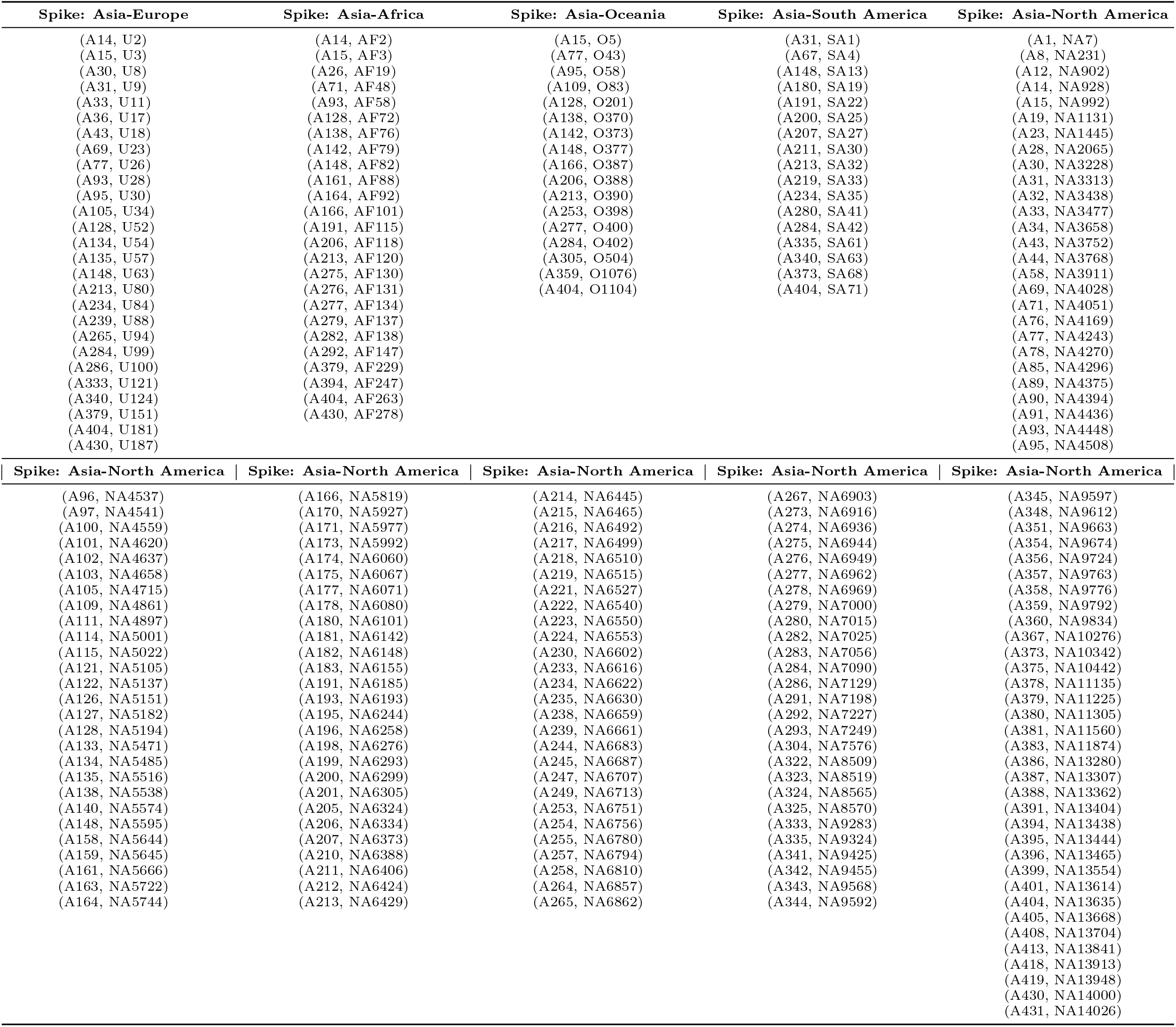
List of pairs of identical spike proteins of SARS-CoV-2 originated from six continents

**Table 10:**
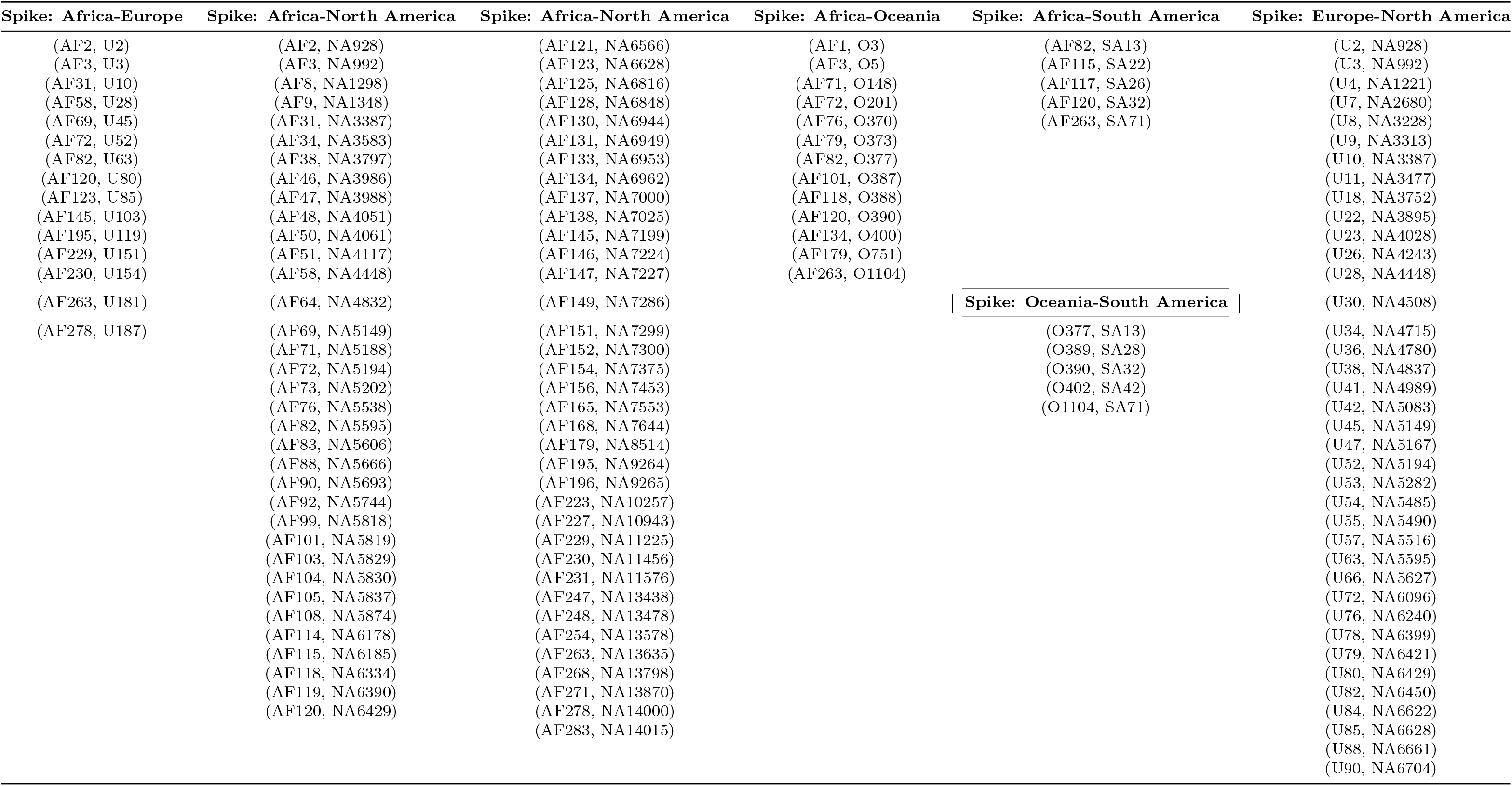
List of pairs of identical spike proteins of SARS-CoV-2 originated from different continents

**Table 11:**
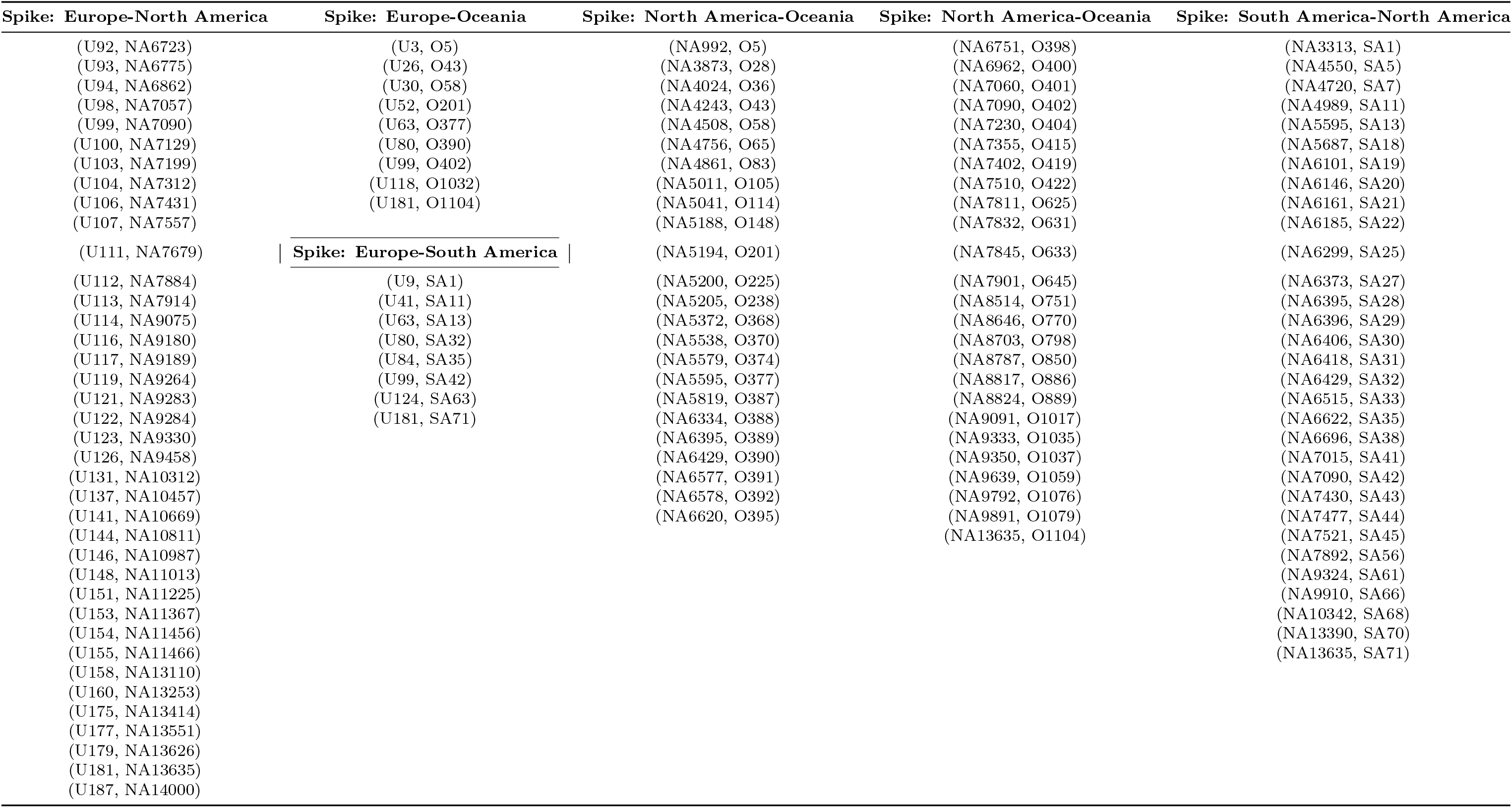
List of pairs of identical spike proteins of SARS-CoV-2 originated from different continents

**Table 12:** List of spike proteins from Asia, which were found to be identical with spike proteins from other five continents

| *Spike proteins (Asia) which were found to be identical with spike proteins from other five continents* |  |  |  |  |  |  |  |  |
| --- | --- | --- | --- | --- | --- | --- | --- | --- |
| A1 | A71 | A115 | A171 | A207 | A239 | A280 | A344 | A388 |
| A8 | A76 | A121 | A173 | A210 | A244 | A282 | A345 | A391 |
| A12 | A77 | A122 | A174 | A211 | A245 | A283 | A348 | A394 |
| A14 | A78 | A126 | A175 | A212 | A247 | A284 | A351 | A395 |
| A15 | A85 | A127 | A177 | A213 | A249 | A286 | A354 | A396 |
| A19 | A89 | A128 | A178 | A214 | A253 | A291 | A356 | A399 |
| A23 | A90 | A133 | A180 | A215 | A254 | A292 | A357 | A401 |
| A26 | A91 | A134 | A181 | A216 | A255 | A293 | A358 | A404 |
| A28 | A93 | A135 | A182 | A217 | A257 | A304 | A359 | A405 |
| A30 | A95 | A138 | A183 | A218 | A258 | A305 | A360 | A408 |
| A31 | A96 | A140 | A191 | A219 | A264 | A322 | A367 | A413 |
| A32 | A97 | A142 | A193 | A221 | A265 | A323 | A373 | A418 |
| A33 | A100 | A148 | A195 | A222 | A267 | A324 | A375 | A419 |
| A34 | A101 | A158 | A196 | A223 | A273 | A325 | A378 | A430 |
| A36 | A102 | A159 | A198 | A224 | A274 | A333 | A379 | A431 |
| A43 | A103 | A161 | A199 | A230 | A275 | A335 | A380 |  |
| A44 | A105 | A163 | A200 | A233 | A276 | A340 | A381 |  |
| A58 | A109 | A164 | A201 | A234 | A277 | A341 | A383 |  |
| A67 | A111 | A166 | A205 | A235 | A278 | A342 | A386 |  |
| A69 | A114 | A170 | A206 | A238 | A279 | A343 | A387 |  |

**Table 13:** List of spike proteins from Africa, which were found to be identical with spike proteins from other five continents

| *Spike proteins (Africa) which were found to be identical with spike proteins from other five continents* |  |  |  |  |  |  |  |  |  |  |
| --- | --- | --- | --- | --- | --- | --- | --- | --- | --- | --- |
| AF1 | AF34 | AF58 | AF79 | AF101 | AF117 | AF128 | AF145 | AF156 | AF227 | AF263 |
| AF2 | AF38 | AF64 | AF82 | AF103 | AF118 | AF130 | AF146 | AF165 | AF229 | AF268 |
| AF3 | AF46 | AF69 | AF83 | AF104 | AF119 | AF131 | AF147 | AF168 | AF230 | AF271 |
| AF8 | AF47 | AF71 | AF88 | AF105 | AF120 | AF133 | AF149 | AF179 | AF231 | AF278 |
| AF9 | AF48 | AF72 | AF90 | AF108 | AF121 | AF134 | AF151 | AF195 | AF247 | AF283 |
| AF19 | AF50 | AF73 | AF92 | AF114 | AF123 | AF137 | AF152 | AF196 | AF248 |  |
| AF31 | AF51 | AF76 | AF99 | AF115 | AF125 | AF138 | AF154 | AF223 | AF254 |  |

**Table 14:**
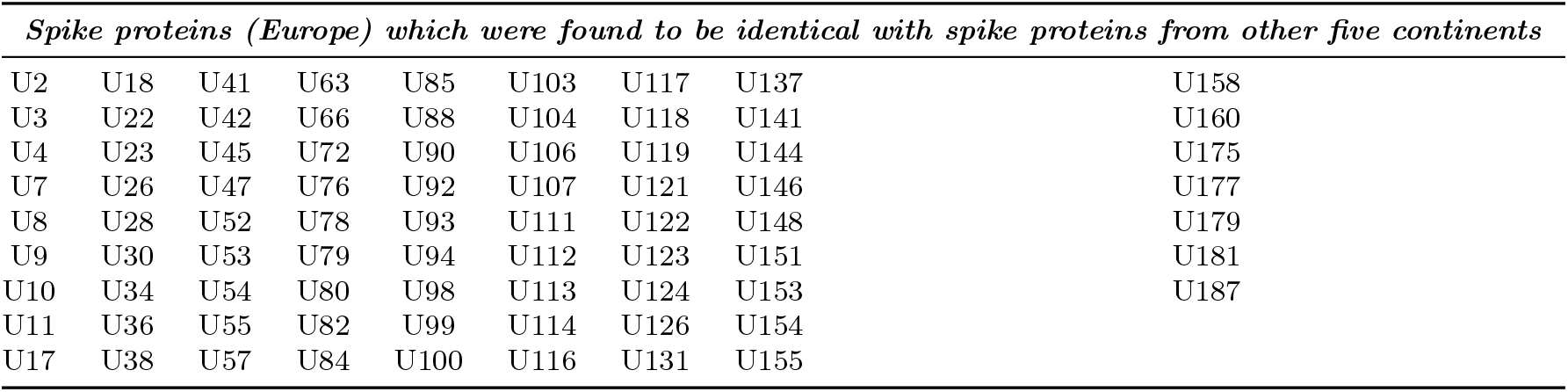
List of spike proteins from Europe, which were found to be identical with spike proteins from other five continents

| *Spike proteins (Europe) which were found to be identical with spike proteins from other five continents* |  |  |  |  |  |  |  |  |
| --- | --- | --- | --- | --- | --- | --- | --- | --- |
| U2 | U18 | U41 | U63 | U85 | U103 | U117 | U137 | U158 |
| U3 | U22 | U42 | U66 | U88 | U104 | U118 | U141 | U160 |
| U4 | U23 | U45 | U72 | U90 | U106 | U119 | U144 | U175 |
| U7 | U26 | U47 | U76 | U92 | U107 | U121 | U146 | U177 |
| U8 | U28 | U52 | U78 | U93 | U111 | U122 | U148 | U179 |
| U9 | U30 | U53 | U79 | U94 | U112 | U123 | U151 | U181 |
| U10 | U34 | U54 | U80 | U98 | U113 | U124 | U153 | U187 |
| U11 | U36 | U55 | U82 | U99 | U114 | U126 | U154 |  |
| U17 | U38 | U57 | U84 | U100 | U116 | U131 | U155 |  |

**Table 15:** List of spike proteins from North America, which were found to be identical with spike proteins from other five continents

| *Spike proteins (North America) which were found to be identical with spike proteins from other five continents* |  |  |  |  |  |  |  |  |  |  |
| --- | --- | --- | --- | --- | --- | --- | --- | --- | --- | --- |
| NA7 | NA3911 | NA4837 | NA5595 | NA6161 | NA6510 | NA6810 | NA7300 | NA8703 | NA9792 | NA13390 |
| NA231 | NA3986 | NA4861 | NA5606 | NA6178 | NA6515 | NA6816 | NA7312 | NA8787 | NA9834 | NA13404 |
| NA377 | NA3988 | NA4897 | NA5627 | NA6185 | NA6527 | NA6848 | NA7355 | NA8817 | NA9891 | NA13414 |
| NA389 | NA4024 | NA4989 | NA5644 | NA6193 | NA6540 | NA6857 | NA7375 | NA8824 | NA9910 | NA13438 |
| NA390 | NA4028 | NA5001 | NA5645 | NA6240 | NA6550 | NA6862 | NA7402 | NA9075 | NA10257 | NA13444 |
| NA402 | NA4051 | NA5011 | NA5666 | NA6244 | NA6553 | NA6903 | NA7430 | NA9091 | NA10276 | NA13465 |
| NA902 | NA4061 | NA5022 | NA5687 | NA6258 | NA6566 | NA6916 | NA7431 | NA9180 | NA10312 | NA13478 |
| NA928 | NA4117 | NA5041 | NA5693 | NA6276 | NA6577 | NA6936 | NA7453 | NA9189 | NA10342 | NA13551 |
| NA992 | NA4169 | NA5083 | NA5722 | NA6293 | NA6578 | NA6944 | NA7477 | NA9264 | NA10442 | NA13554 |
| NA1104 | NA4243 | NA5105 | NA5744 | NA6299 | NA6602 | NA6949 | NA7510 | NA9265 | NA10457 | NA13578 |
| NA1131 | NA4270 | NA5137 | NA5818 | NA6305 | NA6616 | NA6953 | NA7521 | NA9283 | NA10669 | NA13614 |
| NA1221 | NA4296 | NA5149 | NA5819 | NA6324 | NA6620 | NA6962 | NA7553 | NA9284 | NA10811 | NA13626 |
| NA1298 | NA4375 | NA5151 | NA5829 | NA6334 | NA6622 | NA6969 | NA7557 | NA9324 | NA10943 | NA13635 |
| NA1348 | NA4394 | NA5167 | NA5830 | NA6373 | NA6628 | NA7000 | NA7576 | NA9330 | NA10987 | NA13668 |
| NA1445 | NA4436 | NA5182 | NA5837 | NA6388 | NA6630 | NA7015 | NA7644 | NA9333 | NA11013 | NA13704 |
| NA2065 | NA4448 | NA5188 | NA5874 | NA6390 | NA6659 | NA7025 | NA7679 | NA9350 | NA11135 | NA13798 |
| NA2680 | NA4508 | NA5194 | NA5927 | NA6395 | NA6661 | NA7056 | NA7811 | NA9425 | NA11225 | NA13841 |
| NA3228 | NA4537 | NA5200 | NA5977 | NA6396 | NA6683 | NA7057 | NA7832 | NA9455 | NA11305 | NA13870 |
| NA3313 | NA4541 | NA5202 | NA5992 | NA6399 | NA6687 | NA7060 | NA7845 | NA9458 | NA11367 | NA13913 |
| NA3387 | NA4550 | NA5205 | NA6060 | NA6406 | NA6696 | NA7090 | NA7884 | NA9568 | NA11456 | NA13948 |
| NA3438 | NA4559 | NA5282 | NA6067 | NA6418 | NA6704 | NA7129 | NA7892 | NA9592 | NA11466 | NA14000 |
| NA3477 | NA4620 | NA5372 | NA6071 | NA6421 | NA6707 | NA7198 | NA7901 | NA9597 | NA11560 | NA14015 |
| NA3583 | NA4637 | NA5471 | NA6080 | NA6424 | NA6713 | NA7199 | NA7914 | NA9612 | NA11576 | NA14026 |
| NA3658 | NA4658 | NA5485 | NA6096 | NA6429 | NA6723 | NA7224 | NA8509 | NA9639 | NA11874 |  |
| NA3752 | NA4715 | NA5490 | NA6101 | NA6445 | NA6751 | NA7227 | NA8514 | NA9663 | NA13110 |  |
| NA3768 | NA4720 | NA5516 | NA6142 | NA6450 | NA6756 | NA7230 | NA8519 | NA9674 | NA13253 |  |
| NA3797 | NA4756 | NA5538 | NA6146 | NA6465 | NA6775 | NA7249 | NA8565 | NA9724 | NA13280 |  |
| NA3873 | NA4780 | NA5574 | NA6148 | NA6492 | NA6780 | NA7286 | NA8570 | NA9763 | NA13307 |  |
| NA3895 | NA4832 | NA5579 | NA6155 | NA6499 | NA6794 | NA7299 | NA8646 | NA9776 | NA13362 |  |

**Table 16:** List of spike proteins from Oceania, which were found to be identical with spike proteins from other five continents

| *Spike proteins (Oceania) which were found to be identical with spike proteins from other five continents* |  |  |  |  |  |  |  |  |  |
| --- | --- | --- | --- | --- | --- | --- | --- | --- | --- |
| O3 | O105 | O373 | O392 | O419 | O770 |  |  |  | O1037 |
| O5 | O114 | O374 | O395 | O422 | O798 |  |  |  | O1059 |
| O28 | O148 | O377 | O398 | O504 | O850 |  |  |  | O1076 |
| O36 | O201 | O387 | O400 | O625 | O886 |  |  |  | O1079 |
| O43 | O225 | O388 | O401 | O631 | O889 |  |  |  | O1104 |
| O58 | O238 | O389 | O402 | O633 | O1017 |  |  |  |  |
| O65 | O368 | O390 | O404 | O645 | O1032 |  |  |  |  |
| O83 | O370 | O391 | O415 | O751 | O1035 |  |  |  |  |

**Table 17:** List of spike proteins from South America, which were found to be identical with spike proteins from other five continents

| *Spike proteins (Oceania) which were found to be identical with spike proteins from other five continents* |  |  |  |  |  |  |  |  |  |
| --- | --- | --- | --- | --- | --- | --- | --- | --- | --- |
| SA1 | SA13 | SA22 | SA29 | SA35 | SA44 |  |  |  | SA66 |
| SA4 | SA18 | SA25 | SA30 | SA38 | SA45 |  |  |  | SA68 |
| SA5 | SA19 | SA26 | SA31 | SA41 | SA56 |  |  |  | SA70 |
| SA7 | SA20 | SA27 | SA32 | SA42 | SA61 |  |  |  | SA71 |
| SA11 | SA21 | SA28 | SA33 | SA43 | SA63 |  |  |  |  |

It was noticed that IPs for all the unique S proteins from the six continents were distributed in between 5.61 and 7.79. The largest interval of IPs was found for the unique S proteins from North America. Therefore, the widest varieties of unique S proteins were found in North America.

The degree of non-linearity of the plots of IPs for each protein from each continent shows wide variations of unique S proteins (Figures 7 and 8).

**Figure 7:**
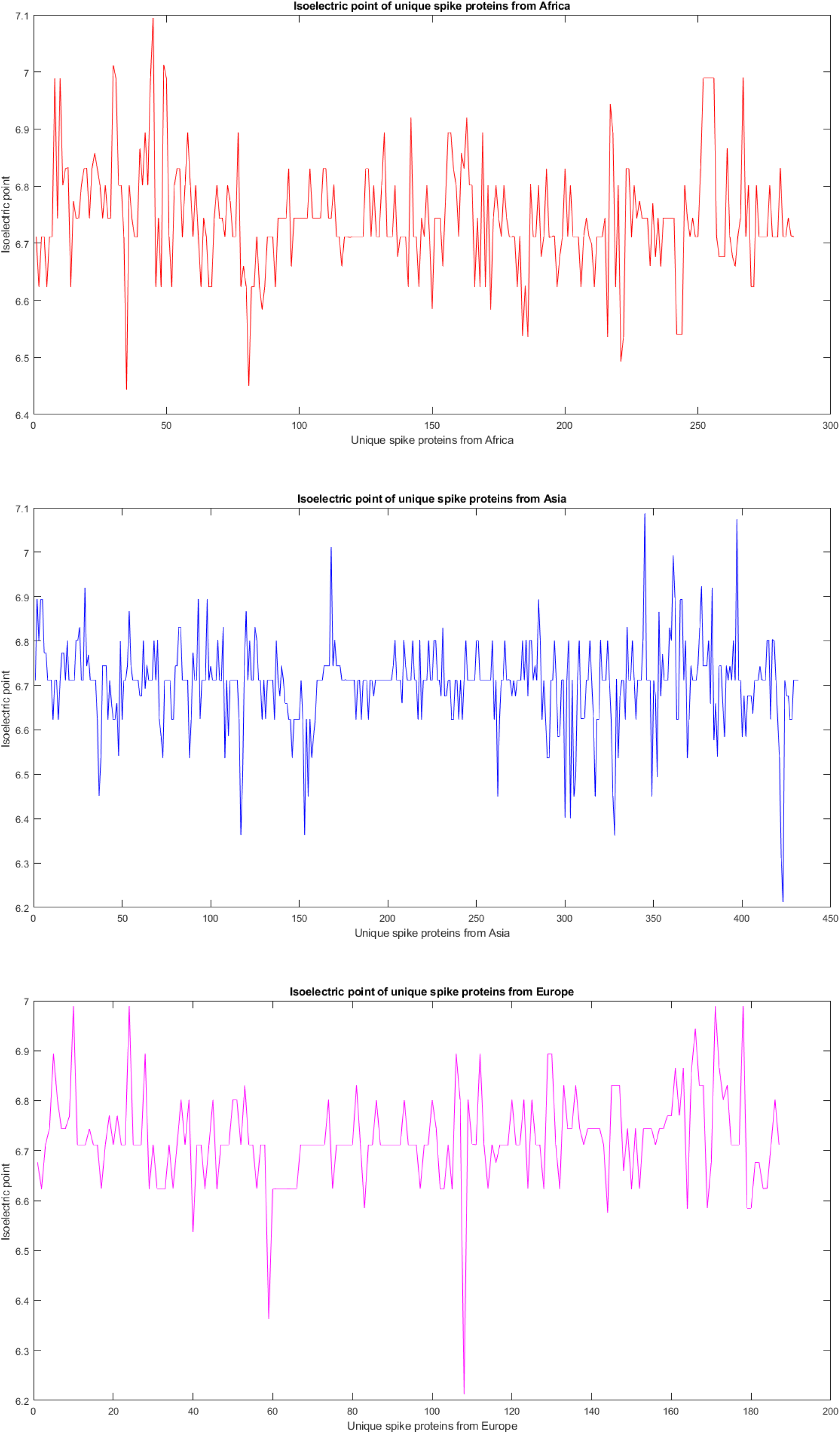
Isoelectric point of unique S proteins from different continents

**Figure 8:**
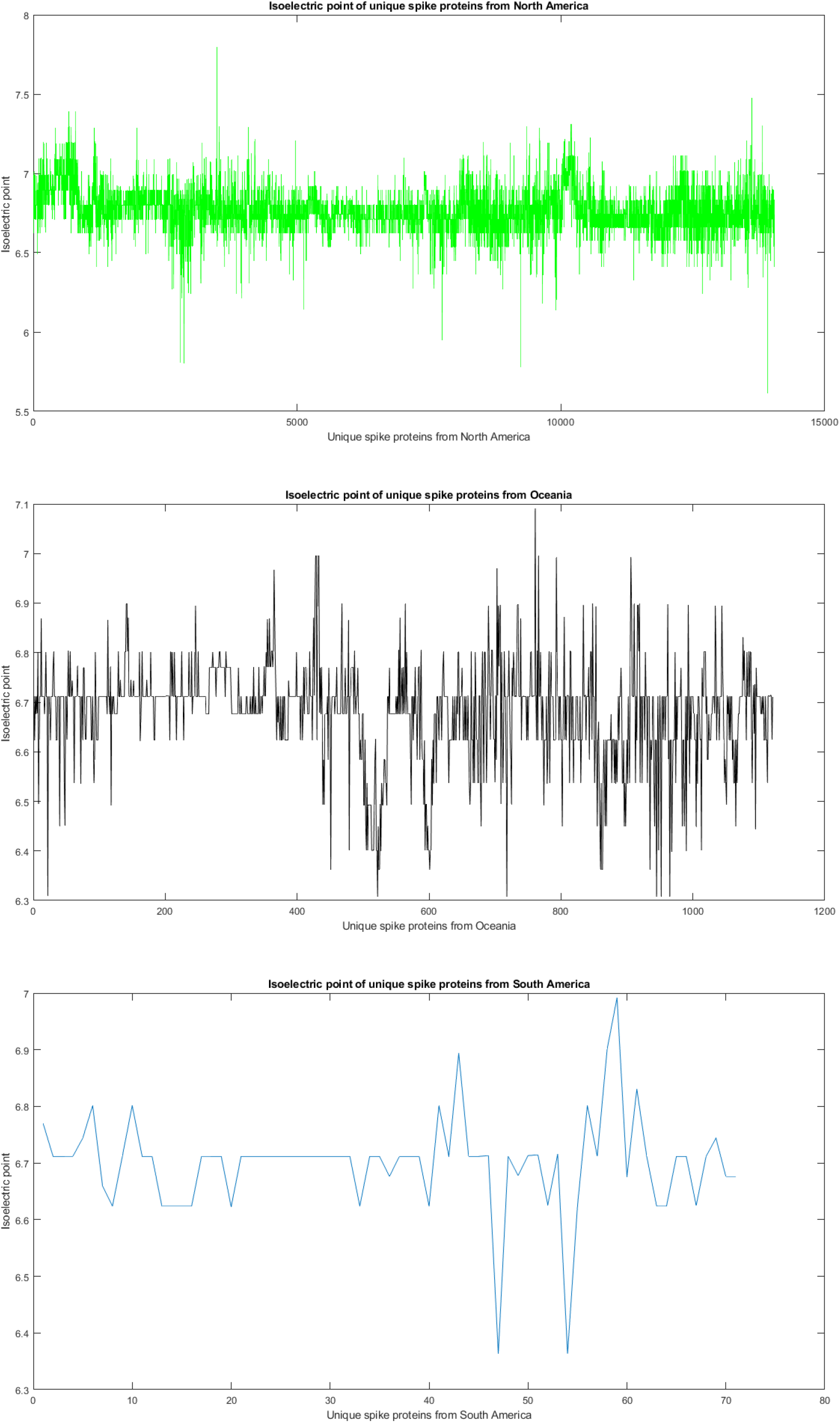
Isoelectric point of unique S proteins from different continents

## 4. Discussion and concluding remarks

Various mutations in S proteins lead to the evolution of new variants of SARS-CoV-2 [29]. Naturally, our attention was captured to characterize unique S protein variants which were embedded in SARS-CoV-2 genomes infecting millions people worldwide [30]. As of May 7, 2021, there are 127760 patients infected with SARS-CoV-2 with 16143 S protein variants, which undoubtedly well-organized by means of amino acids composition and conservation as it was depicted by Shannon entropy and isoelectric point. Among the unique spike proteins present in a continent, many of them are common in other continents as well (Table 2). On the other hand, there is still a handful of unique spike protein variants residing in each continent. Considering the nature and biological implications of the new variants of SARS-CoV-2 caused by different mutations in S proteins, the appearance of several unique S variants in SARS-CoV-2 is certainly an worrying event. [31]. There are still many unique S protein variants in all continents that may spread from person to person through close communities or by spontaneous mutations caused a condition that may become alarming.

We observed that unique S proteins from North America have mutations in almost every amino acid residue position (1184 out of 1273), while unique spike variants from the other continents only have mutations in 16 to 20% of residues. So, even if international travel is limited, S proteins from these five continents will likely acquire mutations at other residue positions where mutations have already been found in the specific variants from North America due to natural evolution. Based on the amino acid frequency distributions in the S protein variants from all the continents, a phylogenetic relationship among the continents was drawn. The phylogenetic relationship implies that unique S proteins from North America were found to be significantly different from that of other five continents. Therefore, the possibility of spreading the unique variants originated from North America to the other geographic locations by means of international travel is high, and numerous mutations have been detected already in the unique variants from North America. Of note, South America infection/herd immunity status may have summarized by Manaus city example (the capital of Amazonas state in northern Brazil) where by June 2020 to October 2020 SARS-CoV-2 prevalence among Manaus’ population increased from ¿60% to ¿70%, a condition which may mirror acquisition of herd immunity [32]. By January 2021 Manaus had a huge resurgence in cases due to emergence of a new variant known as P.1 which was responsible for nearly 100% of the new case [33]. Although the population may have then reached a high herd immunity threshold, there is still a risk of resurgence of new immunity-escape variants, which raises important questions. For example, Is post-infection herd immunity not enough for protection and should it be combined with vaccination? 2. Will the crucial viral variants (mutations) be listed by WHO and recommended to be included in “next generation vaccines”? [34, 35]. In addition, we cannot yet exclude the possibility of serious mutations in the viral RBD emerging in India and the USA [34].

Hence in the near future, we can expect to experience more new SARS-CoV-2 variants which might cause third, fourth, and fifth etc. waves of COVID-19. Therefore, massive vaccination is necessary to combat COVID-19, and of course, existing vaccines must be reviewed, and if needed further re-engineered may be required based on newly emerging S protein variants.

## Acknowledgement

SSH devised the study. SSH and VNU contributed to the implementation of the research, to the analysis of the results and to the writing of the initial draft of the manuscript. KL, PPC, BDU, RK, AB, MS, AL, SPS, GKA, AAAA, ASA, PA, TMAEA, EMR, and KT reviewed and edited the manuscript. NR and MT proof-read the manuscript. All authors read the final version of the manuscript and approve.

## Conflict of interests statement

Authors have no conflict of interest to declare.

## Notes

### Competing Interest Statement

The authors have declared no competing interest.

## References

[1] S. M. Lokman, M. Rasheduzzaman, A. Salauddin, R. Barua, A. Y. Tanzina, M. H. Rumi, M. I. Hossain, A. Z. Siddiki, A. Mannan, M. M. Hasan, Exploring the genomic and proteomic variations of sars-cov-2 spike glycoprotein: a computational biology approach, Infection, Genetics and Evolution 84 (2020) 104389.

[2] A. Serrano-Aroca, K. Takayama, A. Tuñón-Molina, M. Seyran, S. S. Hassan, P. P. Choudhury, V. N. Uversky, K. Lundstrom, P. Adadi, G. Palù, et al., Carbon-based nanomaterials: Promising antiviral agents to combat covid-19 in the microbial resistant era, ACS Nano PMID: 33826850. doi: 10.1021/acsnano.1c00629.

[3] S. Hassan, S. Ghosh, D. Attrish, P. P. Choudhury, A. A. Aljabali, B. D. Uhal, K. Lundstrom, N. Rezaei, V. N. Uversky, M. Seyran, et al., Possible transmission flow of sars-cov-2 based on ace2 features, Molecules 25 (24) (2020) 5906.

[4] M. Martí, A. Tuñón-Molina, F. L. Aachmann, Y. Muramoto, T. Noda, K. Takayama, A. Serrano-Aroca, Protective face mask filter capable of inactivating sars-cov-2, and methicillin-resistant staphylococcus aureus and staphylococcus epidermidis, Polymers 13 (2) (2021) 207.

[5] S. S. Hassan, D. Attrish, S. Ghosh, P. P. Choudhury, V. N. Uversky, A. A. Aljabali, K. Lundstrom, B. D. Uhal, N. Rezaei, M. Seyran, et al., Notable sequence homology of the orf10 protein introspects the architecture of sars-cov-2, International Journal of Biological Macromolecules 181 (2021) 801–809.

[6] S. S. Hassan, A. A. Aljabali, P. K. Panda, S. Ghosh, D. Attrish, P. P. Choudhury, M. Seyran, D. Pizzol, P. Adadi, T. M. Abd El-Aziz, et al., A unique view of sars-cov-2 through the lens of orf8 protein, Computers in biology and medicine (2021) 104380.

[7] L. Zhang, C. B. Jackson, H. Mou, A. Ojha, H. Peng, B. D. Quinlan, E. S. Rangarajan, A. Pan, A. Vanderheiden, M. S. Suthar, et al., Sars-cov-2 spike-protein d614g mutation increases virion spike density and infectivity, Nature communications 11 (1) (2020) 1–9.

[8] L. Guruprasad, Human sars cov-2 spike protein mutations, Proteins: Structure, Function, and Bioinformatics 89 (5) (2021) 569–576.

[9] R. Henderson, R. J. Edwards, K. Mansouri, K. Janowska, V. Stalls, S. M. Gobeil, M. Kopp, D. Li, R. Parks, A. L. Hsu, et al., Controlling the sars-cov-2 spike glycoprotein conformation, Nature structural & molecular biology 27 (10) (2020) 925–933.

[10] M. Seyran, K. Takayama, V. N. Uversky, K. Lundstrom, G. Palù, S. P. Sherchan, D. Attrish, N. Rezaei, A. A. Aljabali, S. Ghosh, et al., The structural basis of accelerated host cell entry by sars-cov-2, The FEBS journal (2020).

[11] E. B. Hodcroft, D. B. Domman, D. J. Snyder, K. Oguntuyo, M. Van Diest, K. H. Densmore, K. C. Schwalm, J. Femling, J. L. Carroll, R. S. Scott, et al., Emergence in late 2020 of multiple lineages of sars-cov-2 spike protein variants affecting amino acid position 677, MedRxiv (2021).

[12] Z. Ke, J. Oton, K. Qu, M. Cortese, V. Zila, L. McKeane, T. Nakane, J. Zivanov, C. J. Neufeldt, B. Cerikan, et al., Structures and distributions of sars-cov-2 spike proteins on intact virions, Nature 588 (7838) (2020) 498–502.

[13] O. A. MacLean, R. J. Orton, J. B. Singer, D. L. Robertson, No evidence for distinct types in the evolution of sars-cov-2, Virus Evolution 6 (1) (2020) veaa034.

[14] L. van Dorp, M. Acman, D. Richard, L. P. Shaw, C. E. Ford, L. Ormond, C. J. Owen, J. Pang, C. C. Tan, F. A. Boshier, et al., Emergence of genomic diversity and recurrent mutations in sars-cov-2, Infection, Genetics and Evolution 83 (2020) 104351.

[15] J. Zhang, Y. Cai, T. Xiao, J. Lu, H. Peng, S. M. Sterling, R. M. Walsh, S. Rits-Volloch, H. Zhu, A. N. Woosley, et al., Structural impact on sars-cov-2 spike protein by d614g substitution, Science 372 (6541) (2021) 525–530.

[16] S. E. Park, Epidemiology, virology, and clinical features of severe acute respiratory syndrome-coronavirus-2 (sars-cov-2; coronavirus disease-19), Clinical and experimental pediatrics 63 (4) (2020) 119.

[17] E. Callaway, The coronavirus is mutating-does it matter?, Nature 585 (7824) (2020) 174–177.

[18] B. Korber, W. M. Fischer, S. Gnanakaran, H. Yoon, J. Theiler, W. Abfalterer, N. Hengartner, E. E. Giorgi, T. Bhattacharya, B. Foley, et al., Tracking changes in sars-cov-2 spike: evidence that d614g increases infectivity of the covid-19 virus, Cell 182 (4) (2020) 812–827.

[19] E. Volz, V. Hill, J. T. McCrone, A. Price, D. Jorgensen, Á. O’Toole, J. Southgate, R. Johnson, B. Jackson, F. F. Nascimento, et al., Evaluating the effects of sars-cov-2 spike mutation d614g on transmissibility and pathogenicity, Cell 184 (1) (2021) 64–75.

[20] T. C. Williams, W. A. Burgers, Sars-cov-2 evolution and vaccines: cause for concern?, The Lancet Respiratory Medicine 9 (4) (2021) 333–335.

[21] H. Tegally, E. Wilkinson, M. Giovanetti, A. Iranzadeh, V. Fonseca, J. Giandhari, D. Doolabh, S. Pillay, E. J. San, N. Msomi, et al., Emergence of a sars-cov-2 variant of concern with mutations in spike glycoprotein., Nature (2021).

[22] D. J. Brooks, J. R. Fresco, A. M. Lesk, M. Singh, Evolution of amino acid frequencies in proteins over deep time: inferred order of introduction of amino acids into the genetic code, Molecular Biology and Evolution 19 (10) (2002) 1645–1655.

[23] V. Vacic, V. N. Uversky, A. K. Dunker, S. Lonardi, Composition profiler: a tool for discovery and visualization of amino acid composition differences, BMC bioinformatics 8 (1) (2007) 1–7.

[24] M. Sickmeier, J. A. Hamilton, T. LeGall, V. Vacic, M. S. Cortese, A. Tantos, B. Szabo, P. Tompa, J. Chen, V. N. Uversky, et al., Disprot: the database of disordered proteins, Nucleic acids research 35 (suppl_1) (2007) D786–D793.

[25] S. S. Hassan, D. Attrish, S. Ghosh, P. P. Choudhury, B. Roy, Pathogenetic perspective of missense mutations of orf3a protein of sars-cov-2, Virus Research (2021) 198441.

[26] S. S. Hassan, P. P. Choudhury, B. Roy, S. S. Jana, Missense mutations in sars-cov2 genomes from indian patients, Genomics 112 (6) (2020) 4622–4627.

[27] B. J. Strait, T. G. Dewey, The shannon information entropy of protein sequences, Biophysical journal 71 (1) (1996) 148–155.

[28] P. G. Righetti, Determination of the isoelectric point of proteins by capillary isoelectric focusing, Journal of chromatography A 1037 (1-2) (2004) 491–499.

[29] A. Baum, B. O. Fulton, E. Wloga, R. Copin, K. E. Pascal, V. Russo, S. Giordano, K. Lanza, N. Negron, M. Ni, et al., Antibody cocktail to sars-cov-2 spike protein prevents rapid mutational escape seen with individual antibodies, Science 369 (6506) (2020) 1014–1018.

[30] Z. Liu, L. A. VanBlargan, L.-M. Bloyet, P. W. Rothlauf, R. E. Chen, S. Stumpf, H. Zhao, J. M. Errico, E. S. Theel, M. J. Liebeskind, et al., Identification of sars-cov-2 spike mutations that attenuate monoclonal and serum antibody neutralization, Cell host & microbe 29 (3) (2021) 477–488.

[31] B. Dearlove, E. Lewitus, H. Bai, Y. Li, D. B. Reeves, M. G. Joyce, P. T. Scott, M. F. Amare, S. Vasan, N. L. Michael, et al., A sars-cov-2 vaccine candidate would likely match all currently circulating variants, Proceedings of the National Academy of Sciences 117 (38) (2020) 23652–23662.

[32] L. F. Buss, C. A. Prete, C. M. Abrahim, A. Mendrone, T. Salomon, C. de Almeida-Neto, R. F. França, M. C. Belotti, M. P. Carvalho, A. G. Costa, et al., Three-quarters attack rate of sars-cov-2 in the brazilian amazon during a largely unmitigated epidemic, Science 371 (6526) (2021) 288–292.

[33] C. Aschwanden, Five reasons why covid herd immunity is probably impossible., Nature 591 (7851) (2021) 520–522.

[34] R. K. Gupta, Will sars-cov-2 variants of concern affect the promise of vaccines?, Nature Reviews Immunology (2021) 1–2.

[35] E. M. Redwan, Covid-19 pandemic and vaccination build herd immunity, Eur Rev Med Pharmacol Sci 25(2) (2021) 577–579.

